# Graphene oxide-quenching-based fluorescence *in situ* hybridization (G-FISH) to detect RNA in tissue: simple and fast tissue RNA diagnostics

**DOI:** 10.1101/143628

**Authors:** Do Won Hwang, Yoo Ri Choi, Dohyun Kim, Hye Yoon Park, Kyu Wan Kim, Mee Young Kim, Chul-Kee Park, Dong Soo Lee

## Abstract

FISH-based RNA detection in paraffin-embedded tissue can be challenging, with complicated procedures producing uncertain results and poor image quality. Here, we developed a robust RNA detection method based on graphene oxide (GO) quenching and recovery of fluorescence in situ hybridization (G-FISH) in formalin-fixed paraffin-embedded (FFPE) tissues. Using G-FISH technique, the long noncoding BC1 RNA, β-actin mRNA, miR-124a and miR-21 could be detected in the cytoplasm of a mouse brain, primary hippocampal neurons, and glioblastoma multiforme tumor tissues, respectively. G-FISH showed the increased BC1 RNA level in individual hippocampal neurons of Alzheimer’s disease brain. The fluorescence recovered by G-FISH correlated highly with the amount of miR-21, as measured by real time RT-PCR. We propose G-FISH as a simple, fast, inexpensive, and sensitive method for RNA detection, with very low background, which could be applied to a variety of researches or diagnostic purposes.

---

Coding and noncoding RNA genes regulate cellular phenotypes distinctly in multicellular organisms, and their differential expression characterizes the phenotypes of individual cells in tissues, revealing significant spatial heterogeneity and complexity. Fluorescence *in situ* hybridization (FISH) methods enable the visualization of the subtleties of RNA expression that contributes to cellular developmental or pathological changes at the subcellular or organismal scales. Thus, FISH is used routinely to identify disease biomarkers using formalin fixed, paraffin embedded (FFPE) tissue specimens archived for future medical research after long-term storage^[1,2]^. Numerous efforts have been made to improve the image quality of *in situ* RNA detection, especially for microRNAs (miRNAs), as well as for mRNAs, at subcellular resolution^[3–5]^. However, the intricate and laborious conventional FISH methods for FFPE tissue sections and the concomitant poor image quality have prevented the easy determination of the expression signatures of RNA biomarkers of interest in clinical practice. Denaturation of fluorescence-labeled probes and off-target hybridization, even for such short molecules as miRNAs, lead to low sensitivity and high background with low specificity^[6–8]^. Herein, we propose a graphene oxide (GO) quenching-based method, termed G-FISH (GO quenching and recovery of fluorescence *in situ* hybridization) to overcome these shortcomings^[9–11]^. GO was used to quench fluorophores attached to nucleic acid probes and to deliver this fluorophore-labeled nucleic acid-GO complex into cells. This method is simpler and faster than the conventional FISH to detect various RNAs, such as long noncoding RNAs (lncRNAs), miRNAs, and mRNAs in FFPE tissue, as well as frozen tissue or live cells cultured *in vitro.*

The fluorescence of a complex comprising a GO nanosheet and a fluorophore-labeled peptide nucleic acid (PNA) was recovered when incubated with tissue specimens by hybridization of the GO-PNA with endogenous target RNAs in tissues on slides. The complementary PNA probe was constructed according to a previous report^[11]^, where it showed high selectivity and stability in live cells for *in vitro* miRNA detection. We used single fluorescence labeling at first and then quadruple fluorescence dyes coupled to both the 5’ and 3’-ends of the PNA using a short linker. The entire procedure of G-FISH took approximately 3 h, including the procedures of deparaffinization and ethanol serial rehydration. Coupling of the GO nanosheet with the PNA fluorescence probes was fast and stable, and was accomplished by simply mixing them together because of the π-π interaction between GO and the PNA. Hybridization between the PNA and the endogenous target RNAs was also very quick, such that the time to complete the entire procedure was shorter than that of typical FISH methods (Supplementary Fig. 1).

**Figure 1.**
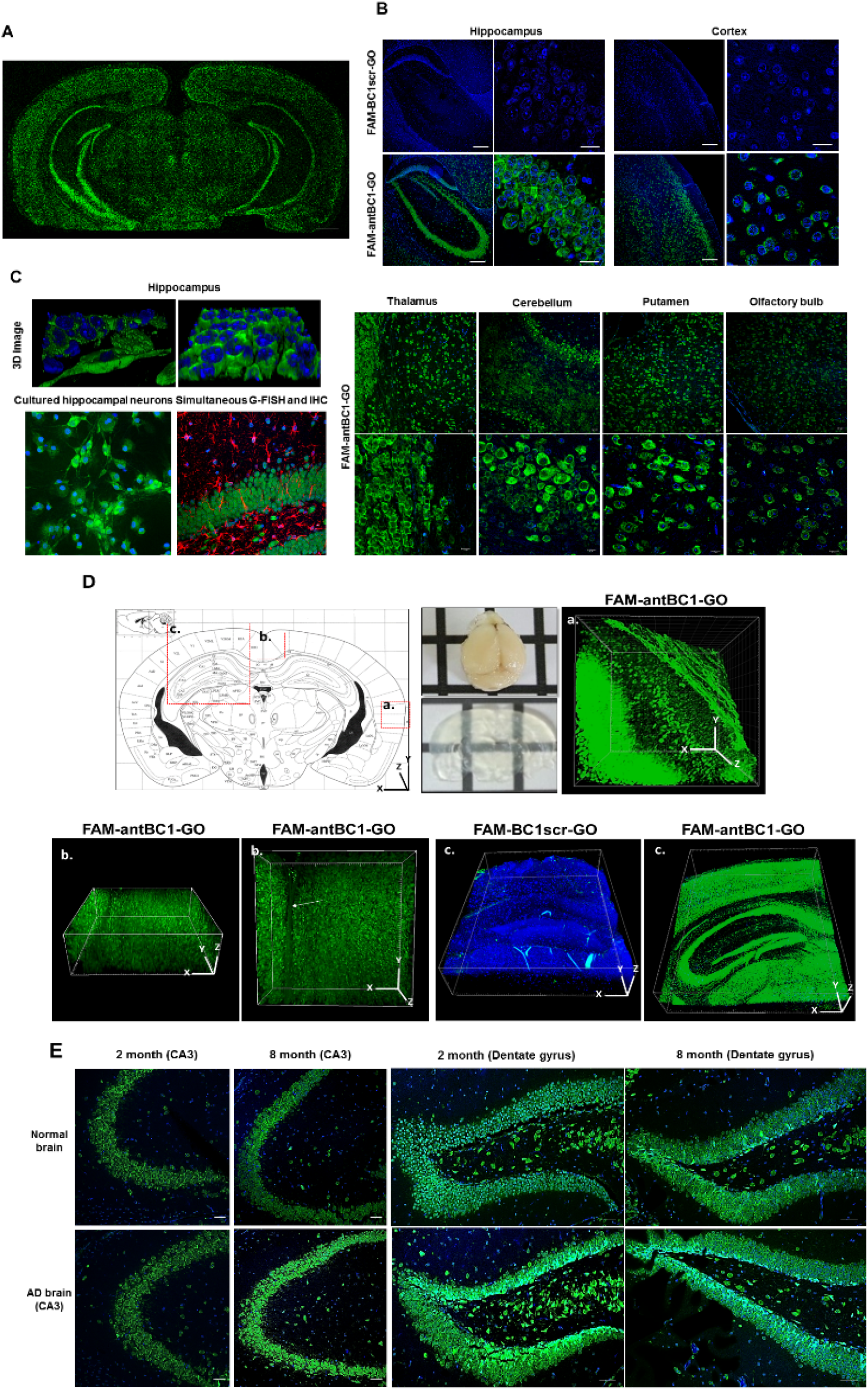
Graphene oxide based quenching of fluorescence *in situ* hybridization (G-FISH) visualization of expression of the long noncoding RNA BC1 in Alzheimer’s disease model mouse brain tissues. (A) Tile scan microscope image showing distribution of BC1 expression on G-FISH on a coronal formalin-fixed paraffin embedded (FFPE) slice of normal mouse brain. (B) Higher magnification of carboxyfluorescein (FAM)-BC1-graphene oxide (GO) showing green fluorescence in the cytoplasm of layered cells in the hippocampus. Left panel: scale bar, 200 μm, right panel: scale bar, 20 μm. Days in (C) Vitro (DIV) 18-dissociated mouse hippocampal neurons were treated with FAM-antBC1-GO. Scale bar, 10 μm. G-FISH for BC1 long noncoding RNA (lncRNA) and immunohistochemistry (IHC) for astrocytes using a glial fibrillary acidic protein (GFAP) antibody were conducted simultaneously using the same brain tissue. Scale bar, 10 μm. (D) Acrylamide hybridized-brain tissues were cleared using an electrophoretic tissue clearing device for lipid removal after the brain tissue was sectioned to 1 mm thickness. These slabs of brain tissue were then incubated with FAM-antBC1-GO or FAM-BC1scr-GO for 16 h in different brain areas (a, b, c) of the mouse brain. (a, b) cortical regions stained with FAM-antBC1-GO, (c) hippocampal region stained with FAM-antBC1-GO (Green), or FAM-BC1scr-GO counterstained with 4’,6-diamidino-2-phenylindole hydrochloride (DAPI) (Blue, 4 × 4 tile scan). (E) Five mutant 5 × familial Alzheimer’s disease (FAD) mice (2 and 8-months old) brains were isolated in an RNA preserving condition. After tissue fixation with 4% formaldehyde and deparaffinization, they were treated with FAM-antBC1-GO.

Simultaneous detection of BC1 long-noncoding RNA and GFAP protein using G-FISH and immunohistochemistry (IHC) with low background in brain tissue. LncRNAs are large and have complicated three-dimensional (3D) structures, with internal sequence complementarity. They also bind to their corresponding RNA binding partner protein in the subcellular environment. This complicated structure of the ribonucleoprotein complex prevents the designing of lncRNA complementary probes with ease using their full-length sequence and requires the exposed open sequence of the lncRNAs to be identified. This is also the case for eukaryotic mRNAs because their long, circular complex chains are organized structurally with eukaryotic translation initiation factors (eIF), ribosomal subunits, and poly A-binding proteins. We addressed this problem by designing the probes using short interfering RNA (siRNA)-based probe screening, as demonstrated previously^[12]^. We first chose a non-translating brain cytoplasmic 1 (BC1) lncRNA that is abundant in the cytoplasm of the neurons of the mouse brain and designed probes specific to BC1^[12]^. Dendritic BC1 lncRNA, which acts as a translational repressor, is associated with a neurological disorder; impairment of BC1 made the mice susceptible to epilepsy^[13]^. The optimal doses of the GO nanosheet and the carboxyfluorescein (FAM)-labeled antisense probe to BC1 (FAM-antBC1) was also determined by fluorescence quenching and de-quenching experiments (Supplementary Fig. 2). Simple treatment with FAM-antBC1-GO nanosheets onto paraffin-embedded coronal brain sections using RNase-free reagents and equipment allowed the visualization of the widespread distribution of BC1 RNA in the tile-scanned coronal image (Fig. 1A). After BC1-GO treatment of the tissue slides, fluorescence started to be recovered within 4 h and reached a maximum at 16 h (Supplementary Fig. 3). The correlation between BC1 expression levels detected by quantitative real-time reverse transcription PCR (qRT-PCR) and G-FISH was high in various brain areas (Supplementary Fig. 4). G-FISH revealed that BC1 RNA signals are mostly localized in the cytoplasmic areas of individual hippocampal and cortical neurons (Fig. 1B). The background fluorescence from non-specific probe binding was minimal, as shown in consecutive coronal slices treated with mismatch-scramble BC1-GO (FAM-BC1scr-GO) (Fig. 1B). BC1 RNA signals were also observed in the thalamus, cerebellum, putamen, and olfactory bulb (Fig. 1C). Interestingly, simultaneous G-FISH and immunohistochemistry (IHC) clearly distinguished astrocytes stained with glial fibrillary acidic protein (GFAP) antibody and neurons visualized by BC1-GO (Fig. 1C, right lower panel). G-FISH revealed BC1 RNA expression in the brain tissue much more clearly than did the conventional FISH method (Supplementary Fig. 5). Treatment with FAM-free antBC1-GO inhibited the interaction of FAM-antBC1-GO with the endogenous BC1 lncRNA competitively (Supplementary Fig. 6). Tissue clearing techniques using **clear lipid-exchanged anatomically rigid imaging/immunostaining-compatible tissue hydrogel** (CLARITY) allow visualization of the elaborate 3D-structural organization of brain slices^[14]^. Combining this technique with G-FISH allowed us to visualize the BC1 expression clearly in the entire 100-μm-thick hippocampus tissues because of the rapid diffusion of the 50–100 nm-sized antBC1-GO nanosheets into the hydrogel-clarified brain section (Fig. 1D). The structural organization of individual pyramidal neurons was observed clearly in the mouse brain. GO can penetrate cells easily through the cell membrane, and thus G-FISH could be applied to live cells with intact membranes in culture^[11,15,16]^. The CLARITY technique allowed antBC1-GO to penetrate to a depth of 100 μm, so that 3D volume imaging of thick brain tissues became possible.

BC200 is a human analog of BC1 and is upregulated markedly in Alzheimer’s disease (AD) patients. This is thought to result from synaptic remodeling via a compensatory mechanism for neuronal degeneration^[17,18]^. BC200 expression in the hippocampus correlated highly with clinical dementia rating scores, suggesting that it could play a role in AD. In this study, we used 5 × familial Alzheimer’s disease (FAD) mice, which have five gene mutations predisposed to an AD-like phenotype, in which we confirmed that they showed spatial memory impairment in the Y-maze test at 8 months of age (Supplementary Fig. 7). Thioflavin-S-stained amyloid deposition was also observed in the cortex, hippocampus, and thalamus at 4 and 8 months of age (Supplementary Fig. 8). In the G-FISH experiment, BC1 expression was increased significantly in CA3 hippocampal neurons of 8-month-old 5 × FAD AD model mice, with rapidly developing severe amyloid plaque pathology, compared with that in age-matched non-AD mice (Fig. 1E). The majority of these neurons with high BC1 expression exhibited an extensive irregular and shrunk cellular shape. Interestingly, in the 5 × FAD mice, the distribution of BC1 in the dentate gyrus appeared to be heterogeneous as early as two months of age, showing intense expression along the subgranular zone where the neural stem cells reside (Fig. 1E). The elevated numbers of BC1 positive cells were positioned mostly around the intergranular layer, showing atypical morphological features, with dystrophic distortion (Fig. 1E). G-FISH could detect the increased BC1 expression that might be associated with impaired neural stem cells in the dentate gyrus.

G-FISH could detect mRNA using a FAM-labeled PNA probe for β-actin mRNA (βAct). Simple treatment with FAM-antβAct-GO on deparaffinized coronal brain tissue sections showed prominent fluorescence of β-actin mRNAs in the cytoplasm of the cells in the hippocampus and cortex (Fig. 2A), while FAM-antAct_scr (mismatched scramble)-GO did not.

**Figure 2.**
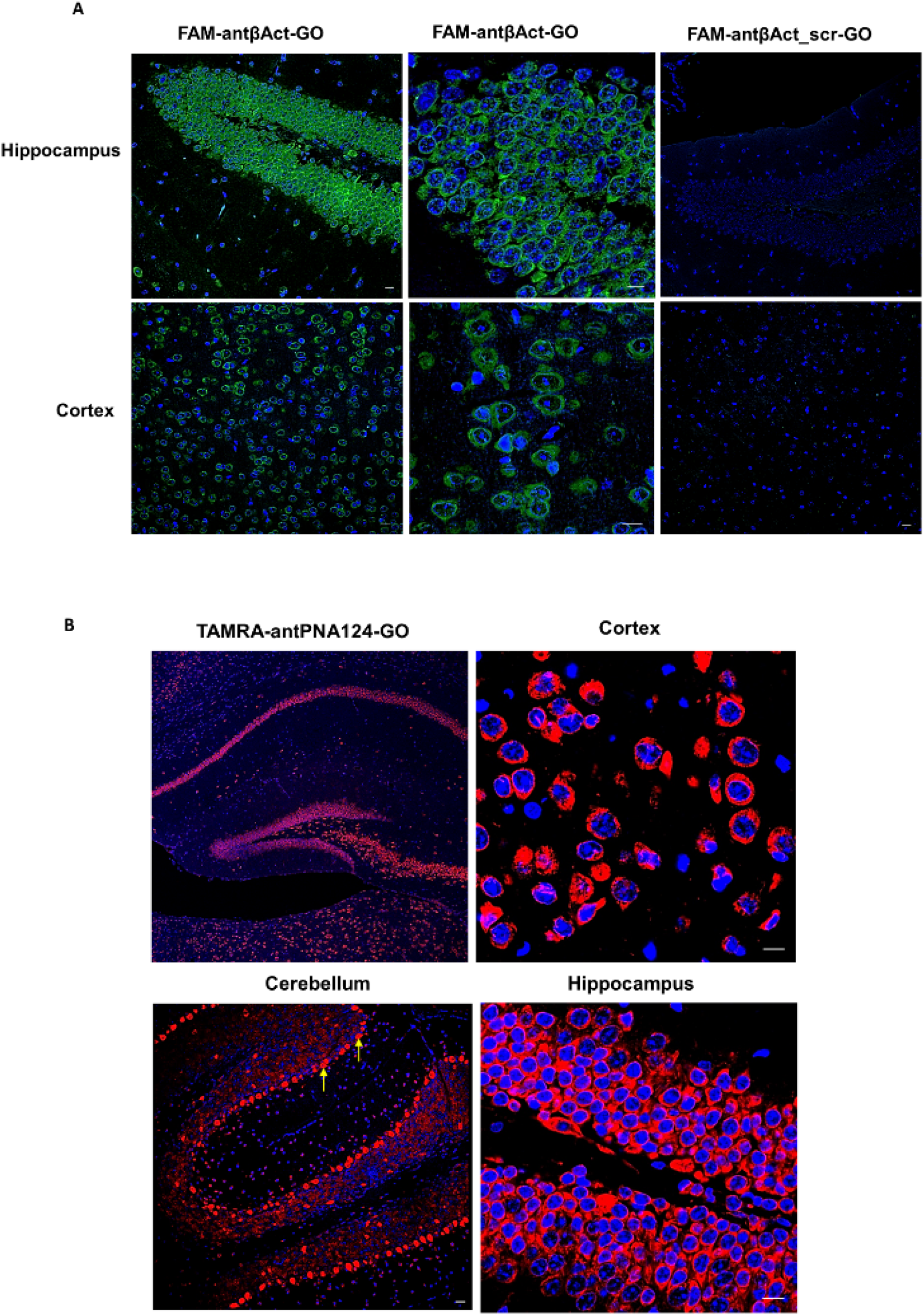
β-actin mRNA and neuron-specific miR-124a detection using G-FISH in brain tissue. (A) FAM-labeled peptide-nucleic acid (PNA) probe for β-actin mRNA (βAct), FAM-antβAct-GO, could visualize constitutive β-actin mRNA in formalin-fixed paraffin embedded (FFPE) coronal brain sections of normal mice. No FAM signals were found using FAM-antβAct_scr-GO. Scale bar, 10 μm. (B) Tetramethylrhodamine (TAMRA)-antPNA124-GO revealed the distribution of miR-124a expression in the cytoplasm of the cortical and hippocampal neurons. Intense TAMRA fluorescence was seen in the Purkinje cell layer (yellow arrow). Scale bar, 20 μm.

G-FISH could also detect the small non-coding miR-124a, which is expressed exclusively in neurons. Tetramethylrhodamine (TAMRA)-antPNA124a-GO showed prominent cytoplasmic fluorescence in hippocampal and cortical neurons (Fig. 2B). The abundant miR-124a expression in Purkinje cells agreed with the observations previous reports using FISH^[4,19]^.

MiR-21 is a common oncogenic miRNA that is expressed predominantly in glioblastoma multiforme (GBM). MiR-21 expression in glioma tissue correlated with pathological grade, which led to the proposal that miR-21 might be used as prognostic marker of GBM^[20,21]^. To increase the signal, we used G-FISH with a quadrupled fluorophore. The FAM4-decorated miR-21 probe amplified the signal and FAM4-antPNA21-GO treatment yielded greater fluorescence in the cytoplasm of cultured miR-21-positive cancer cells than the single fluorophore-linked PNA-GO probe (FAM1-antPNA21-GO, Fig. 3A). FAM-free antPNA21-GO blocked the signal from both FAM1 and FAM4-antPNA21-GO almost completely. When FAM4-antPNA21-GO was used to treat GBM tissue specimens, miR-21 was detected only in GBM tumors and not in the normal brain (Fig. 3B). The fluorescence intensities of FAM4-antPNA21-GO in the GBM tumors of 13 patients correlated highly with the level of miR-21 expression measured by quantitative real-time RT-PCR (Fig. 3C). No G-FISH signal was observed in the cells of FAM4-PNA21_scr-GO treated normal brain and tumor tissues (Supplementary Fig. 9). Quantitative real-time RT-PCR showed higher miR-21 expression in GBM tumor tissue than in normal brain tissue (Supplementary Fig. 10).

**Figure 3.**
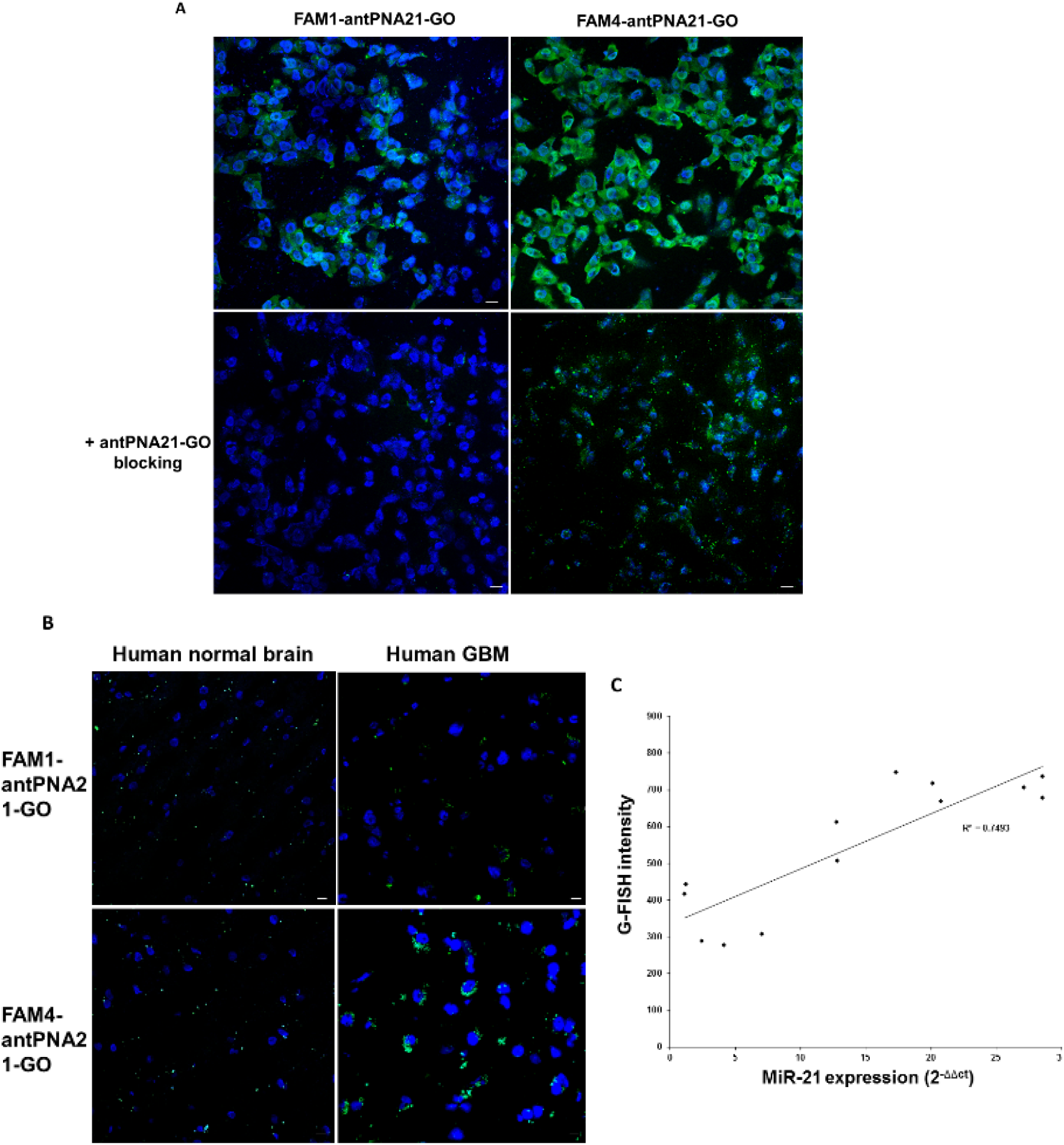
Graphene oxide (GO) based quenching of fluorescence *in situ* hybridization (G-FISH) detection of miR-21 using quadruple carboxyfluorescein (FAM)-PNA21-GO in tissue specimens of human glioblastoma multiforme (GBM) patients. (A) Single and quadruple FAM-antPNA21-GO disclosed miR-21 expression in the cytoplasm of miR-21 positive MDA-MB-231 cells, showing brighter fluorescence with FAM4-antPNA21-GO. Treatment with FAM-free antPNA21-GO blocked the fluorescence recovery of FAM1-or FAM4-antPNA21-GO completely and no signal was found. Scale bar, 20 μm. (B) Single or quadruple FAM1-or-FAM4 antPNA21-GO showed fluorescence in the cytoplasm of cells in glioblastoma multiforme (GBM) tumors, but not in normal human brains. (C) The miR-21 concentration, as measured by quantitative real-time reverse transcription (RT)-PCR, correlated well with the cytoplasmic fluorescence in GBM tumors measured by G-FISH using FAM4-antPNA21-GO

Nanotechnology-based RNA detection in tissue specimens has advanced recently to become faster and more sensitive compared with methods using probes with complicated design and long procedures^[22]^. Probes such as authentic RNAs, locked nucleic acids, PNAs, or even gapmers labeled with various organic fluorophores have been adopted. However, their specificity and high background from off-target binding or incomplete washing have been a serious challenge. In addition, guaranteeing the sensitivity to detect 10 to 20 copies of RNAs per cell has been inhibited by the instability of the probes and hindrance of hybridization by RNA-binding proteins, labeled fluorophores, permeabilizers, temperature, or even cations^[23–26]^. The washing procedure can also cause spurious negative results, even with positive controls using polyd(T) probes or pan-species actin probes, in the case of mRNA detection. G-FISH does not need a washing procedure because the un-hybridized probe attached to GO does not generate non-specific background fluorescence. This was represented in the almost zero background, except for 4’,6-diamidino-2-phenylindole hydrochloride (DAPI)-stained nuclear features, in the specimen. We confirmed that GO in G-FISH was a highly efficient quencher, because mismatched scrambled probes did not yield background fluorescence during BC1, β actin, and miR-124a or miR-21 detection. Hybridization of G-FISH does not require the temperature to be above room temperature (19 °C-21 °C) to detach FAM or TAMRA-labeled PNA probes from GO. De-quenching of the PNA probe can be achieved in mild conditions without many additives, which makes G-FISH simple and robust.

Here, we demonstrated the efficacy of G-FISH by detecting a variety of RNAs with probes constructed using easy and rapid design steps. GO binds readily to single-stranded PNA probes and quenches PNA-linked fluorophores, which increases the sensitivity of RNA detection, producing very low background noise. GO can also penetrate cell membranes and act as a carrier to transport the fluorophore-PNA complex into live cells in culture^[11]^. Deparaffinized FFPE or CLARITY-prepared transparent specimens were penetrable to the fluorophore-PNA-GO. G-FISH using these probes is available widely for many FFPE samples in RNA detection. The G-FISH method can be applied readily to histological specimens of dewaxed FFPE or clarified transparent tissues to better understand RNA expression in order to investigate the meaning of both pathological and physiological processes. Fresh frozen tissues of GBM were also tested as specimens and GO-fluorophore-PNA penetrated well into the cells and visualized the target RNA within the cells.

To assure the integrity of RNA in the specimen of interest, positive controls were included using conventional FISH targeting total mRNAs or constitutively expressing actin or beta-tubulin. We showed that G-FISH could detect beta-actin mRNA expression for quality control of the tissue preparation. The specificity of G-FISH was convincing, especially considering the absolute lack fluorescence using the scramble probes, which maintained the same GC content and length as the target probes. These refinements made the G-FISH experiments much simpler than conventional FISH; G-FISH requires only 3 h for completion of the main procedure and thus is cost-effective.

In summary, G-FISH has the advantage of a very low or no background and thus could be harnessed as a robust diagnostic tool for RNA detection in permanent or frozen tissue samples. It could be used to diagnose diseases, for prognostic studies, and for therapeutic monitoring by preparing already-stored tissues for further clinical retrospective or prospective cohort studies. We use the same preprocessing step of FFPE tissues in G-FISH and conventional FISH; however, the incubation with the G-FISH probes can be performed at room temperature within only 16 h, which will make G-FISH popular for routine use in any laboratory.

Notwithstanding the limitation of G-FISH to obtain single RNA molecule resolution because of the signal on/off switch strategy, G-FISH-based microscopic assessment of the localization and expression signature of an RNA of interest in a heterogeneous cell population will be useful in studies aiming to interpret the complicated molecular heterogeneity of diseased tissues. With advances in visualization tools for microscopy, G-FISH molecular biomarker diagnosis is anticipated. We expect that G-FISH could replace the current FFPE-RNA FISH as a simple and fast RNA detection method for medical diagnostics.

## Experimental

### Section Chemical reaction of GO with FAM-antPNA

The GO nanosheet, synthesized from natural graphite sheets, was purchased from the Lemonex Corporation^[11]^. PNA oligomers were synthesized using solid phase synthesis with Benzothiazole-2-sulfonyl (Bts) pNA monomers, and FAM was labeled on PNA’s 5’-end and Lys’ side chain by an amine-NHS ester reaction (Panagene, Korea). A spacer was inserted to improve labeling reactivity and yield between labeling positions. For the four FAM-PNA, we used the L35 linker (a polyethylene glycol linker) as the spacer.

FAM-antBC1 was reacted with the GO nanosheet a t concentrations ranging from 0.1 μg to 2.0 μg to determine the optimal dose combination. Forty pmoles of FAM-decorated antPNA probes was simply mixed with GO (Tris-HC1, pH 7.4) in an Eppendorf tube and incubated for 10 min at room temperature. FAM-antPNA-GO made with fixed dose of 0.4 μg of GO and 40 pmoles of FAM-antPNA was then reacted with the BC1 oligomer at concentrations from 10 to 80 pmoles at room temperature. The fluorescence intensity was examined quantitatively using GloMax® Discover: Multimode Detection System (Promega Korea, Korea).

### GO quenching of fluorescence *in situ* hybridization (G-FISH) in cultured cells

MDA-MB-231, a human breast cancer cell line, and HeLa, a human cervical cancer cell line, were grown with Dulbecco’s modified Eagle’s medium (DMEM, Welgene, Korea), supplemented with 10% fetal bovine serum (FBS, Invitrogen/LT Korea, Korea), 10 U/ml penicillin, and 10 μg/ml streptomycin at 37 °C in a 5% CO_2_ humidified incubator. Primary hippocampal neurons were isolated from newborn pups of C57BL/6 mice. Hippocampi were dissected, dissociated with trypsin, and plated onto poly-D-lysine coated glass coverslips. The cultures were maintained in Neurobasal-A medium supplemented with B-27, Glutamax, and Primocin (Invivogen/Daeil Bio, Korea) at 37 °C and 5% CO_2_, and then fixed with 4% paraformaldehyde. For the *in vitro* cell study, 1 × 10^5^ cells were seeded on a coverslip in a 24-well plate and treated with pre-made antPNA-GO by mixing 100 pmoles of FAM-PNA and 1 μg of NGO. Fourteen hours after incubation, the cells were fixed with 4% paraformaldehyde and washed twice with 1 × phosphate buffered saline (PBS). For the blocking study, FAM-free antPNA21-GO (PNA complementary to miR-21) and the same amount of FAM-antPNA21-GO, was co-administered to compete for cellular miR-21.

### G-FISH procedures for paraffinized tissue section

The 6-week-old BALB/c male mouse brains were isolated and fixed freshly with 4% paraformaldehyde in PBS, and embedded in paraffin wax. Paraffinized mouse brain tissues were sliced into serial coronal or sagittal sections in RNase-free slides at 4-μm thickness using a microtome. The brain tissue slides from the paraffin embedded brain tissue block were treated with xylene in a Corpin jar for deparaffinization. The slides were treated with a graded ethanol series (50 % diluted in xylene, and 100, 9, 85, 70, and 50 % diluted in distilled water) for rehydration. The samples were rinsed with RNase free water twice. The cells were then immersed in 1× PBS, and then the PBS was wiped out using dust-free tissue paper. FAM-antPNA-GO complexes (1 μg of GO and 100 pmoles of PNA for FAM-antBC1-GO or FAM-antβAct-GO; 5 μg of GO and 500 pmoles of PNA for TAMRA-antPNA124-GO; 9 μg of GO and 300 pmoles of PNA for FAM1-antPNA21-GO; GO: 20 μg, PNA: 300 pmoles for FAM4-antPNA21-GO) were made to a final volume of 100 μl with reaction for 10 min at room temperature (Tris-HCl, pH. 7.4). These complexes were incubated with the slides with mild shaking for 2 h and then further incubated for 16 h in a humidified slide chamber.

### FISH procedure in paraffin brain sectionsre

Mouse brains were fixed with 4% paraformaldehyde in PBS and embedded in paraffin. Paraffinized mouse brains were sectioned at 4-μm thickness using a microtome. Paraffin slice samples were incubated in xylene and an ethanol (100, 95, 70, and 50 % serial rehydration) mixture for 60 min. The samples were digested with 20 μg/mL proteinase K in pre-warmed 50 mM Tris for 20 min at 37 °C for permeabilization. Each slide was rinsed five times in distilled water. Slides were immersed in ice-cold 20 % acetic acid for 20 sec. The slides were dehydrated by washing for approximately 1 min in 70 % ethanol, 95 % ethanol, and 100% ethanol, followed by air-drying. Hybridization solution (100 μL) was added and the slides were incubated for 1 h in a humidified hybridization chamber at 62 °C. After the hybridization solution was removed, 50–100 μL of diluted 5’ digoxygenin-BC1 antisense probe was added to each sample, covering the entire tissue slice. The samples were incubated in the humidified hybridization chamber at 65 °C overnight. Slides were then subjected to the following washes: 50% formamide in 2× saline-sodium citrate (SSC) for 5 min three times at 45 °C; 0.1× SSC 5 min, three times at 65 °C; MABT (maleic acid buffer with tween-20), two times for 30 min at room temperature (RT); and then air dried. Slides were transferred to a humidified chamber and treated with MABT with 2% BSA for 2 h at room temperature. The anti-labeled antibody diluted in MABT with 2% bovine serum albumin (BSA) was incubated for 2 h at room temperature. Slides were then subjected to the following washes: MABT, five times for 10 min; pre-staining buffer (100 mM Tris pH 9.5, 100 mM NaCl, 10 mM MgCl_2_), two times for 10 min; distilled water, two times for 5 min; and then air dried for 30 min. The slides were washed using 100% ethanol, and each sample was mounted with DePeX (Sigma-Aldrich Korea).

### Brain tissue clearing using the CLARITY technique

Adult mice were anesthetized with Avertin (Sigma-Aldrich Korea, Incheon, Korea) and perfused intracardially with cold PBS and 4% paraformaldehyde (PFA). Isolated brains were incubated in 4% PFA for 24 h and transferred into hydrogel monomer solution (4% acrylamide and 0.25% photoinitiator 2,20-Azobis[2-(2-imidazolin-2-yl)propane] dihydrochloride (VA-044, Wako Chemicals, USA) for 24 h at 4 °C. Hydrogel-infused brains were degassed using nitrogen in a vacuum desiccator for 3 min and incubated in a 37 °C water bath for 2–3 h. Excess hydrogel was removed from the polymerized whole brain, which was sliced into 1 mm coronal sections (Roboz Surgical Instrument, USA). The brain sections were transferred into clearing solution (4% SDS in 200 mM boric acid in H_2_O, pH 8.5) in an electrophoresis chamber (Logos Biosystems, Korea), and 45 V was applied at 37 °C. The cleared brains were then washed in PBS overnight and incubated with FAM-antBC1-GO or FAM-BC1scr-NGO, and counterstained with DAPI. After several washing steps, the samples were immersed in CLARITY mounting Solution (RI=1.46, Logos Biosystems, Korea).

### GBM study population and preparation of tissue specimens

Fresh frozen tissue samples were obtained from 13 newly diagnosed supratentorial glioblastoma patients, who were confirmed histologically on surgical resection and biopsy at the Seoul National University Hospital. Tumor tissues were obtained during surgery and then snap-frozen in liquid nitrogen and were stored at −80 °C. Informed consent was obtained from patients before resection, in accordance with the local Institutional Review Board guidelines (IRB No. H 0507-509-153). All the patients were adults with a mean age of 50.9 years (range, 16–82). Tumors were located in the frontal lobe, the temporal lobe, the parietal lobe, or in multiple lobes. In routine diagnostic laboratory analysis for high-grade gliomas, O-6-Methylguanine-DNA Methyltransferase (MGMT) methylation was found in eight patients, and neither the IDH1 mutation nor the 1p/19q co-deletion was observed in any patient.

### Immunofluorescence staining

Paraffinized glass slides were immersed in a fresh solution of xylene, and rehydrated in 100, 95, 90, 80, and 70 % ethanol for 5 min, sequentially. The slides were then rinsed using distilled water and fixed using 0.5 % H_2_O_2_ in methanol. The sections were washed three times in PBS and blocked using goat serum for 30 min. For immunofluorescence, the sections were incubated with primary antibody (GFAP, 1:100, Cell Signaling, USA) diluted in PBS with 5 % BSA in humidified chamber at 4 °C for overnight. The sections were washed using PBS for 5 min twice and then treated with secondary antibody (Alexa Fluor 555 Goat antimouse IgG, 1:100, Life Technologies Korea) in PBS for 1 h at room temperature. After washing with PBS for 5 min, all sections were mounted using Vectashield mounting medium counterstained with 4’,6-diamidino-2-phenylindole hydrochloride (DAPI, Vector Laboratories, UK). Fluorescence images were acquired using a Zeiss LSM 510 META Confocal Imaging System (Carl Zeiss, Korea). The confocal fluorescence filter was set up as follows: For DAPI imaging: excitation/emission wavelength, HFT 358/461 nm; beam splitter pinhole diameter, 98 μm; and filter, BP 420-480 IR. For FAM imaging: excitation/emission wavelength, HFT 494/518 nm; beam splitter pinhole diameter, 84 μm; and filter, BP 505-530 IR. For TAMRA imaging: excitation/emission wavelength, HFT 555/580 nm; filter, BP 560-590 IR.
diameter, 84 μm; and filter, BP 505-530 IR. For TAMRA imaging: excitation/emission wavelength, HFT 555/580 nm; filter, BP 560-590 IR.

### Real time RT-PCR

Total RNA was extracted from human normal brain and human glioblastoma tissue using Trizol (Invitrogen/LT Korea). The purity of total RNA was confirmed using a Nanodrop-1000 Spectrophotometer (Thermo Scientific/LT Korea). Total RNA (2 μg) was transcribed using RnaUsScript reverse transcriptase (LeGene, USA). Reverse transcription (RT)-PCR amplification was performed using an ABI® 7500 instrument (Applied Biosystems, LT Korea) with the SYBR Premix Ex Taq Tli RNaseH Plus (Takara, Korea) SYBR Green Master mix (Clontech Laboratories, Takara, Korea), which is specific for mature mRNA and miRNA sequences. All experiments were conducted in triplicate. RT-Primer listed below. BC1 forward: 5’-TGG GGA TTT AGC TCA GTG GTA G-3’, BC1 reverse: 5’-GTT GTG TGT GCC AGT TAC CTT G-3’, beta-actin forward: 5’-AAG ACC TCT ATG CCA ACA CAG T-3’, beta-actin reverse: 5’-GCT CAG TAA CAG TCC GCC TA-3’, miR-21 (ABM Cat. No. MPH01259), U6 (ABM Cat. No. MPH00001).

### Establishment of 5 × FAD Alzheimer’s disease mouse model

All the animal experimental procedures were approved by Seoul National University Institutional Animal Care and Use Committee and performed in accordance with the guideline from the institute (IACUC: 15-0208-S1A0). The 5 × FAD transgenic mice were generated to overexpress mutant human APP695 with the Swedish (K670N, M671L), Florida (I716V), and London (V717I) mutations, along with human presenilin 1 (PS1) containing two mutations, M146L and L286V. The 2, 4, and 8-month-old 5 × FAD mice performed the Y-maze test and their brains were isolated after intracardiac perfusion.

For habituation to the apparatus of Y-maze test, the mice were placed in the middle of Y-maze and allowed to explore each arm for 24 h before the Y-maze test. The test started in the middle of the Y-maze and the entry frequency into each arm was measured for 8 min. The performance was done in dimmed lighting. The alternation was scored with the pattern of entries without the initial two entries, alternation (%) = number of alternation / total possible alterations × 100. Data were expressed as the means ± SD values. Student’s t-test was used to determine the statistical significance, where p < 0.05 was considered significant.

Isolated brains were placed in 4% PFA for 24 h, and then embedded into paraffin tissue blocks. The paraffin sections were deparaffinized by oven heating and by immersion in xylene and hydrated to distilled water. The hydrated sections were incubated in 1 % Thioflavine S (Santa Cruz Biotechnology, USA) for 5 min and washed in 70 % EtOH for 5 min and in distilled water twice. Fluorescence microscopic observation of mounted tissues was performed using an Olympus BX61 (Olympus, Japan).

### Statistical analysis

Data are displayed as means ± standard deviation (SD) and were tested using Student’s t-test. *P* values of <0.05 were considered statistically significant.

